# Patient-specific haemodynamic simulations of complex aortic dissections informed by commonly available clinical datasets

**DOI:** 10.1101/593780

**Authors:** Mirko Bonfanti, Gaia Franzetti, Gabriele Maritati, Shervanthi Homer-Vanniasinkam, Stavroula Balabani, Vanessa Díaz-Zuccarini

**Author notes:** **Corresponding authors:** E-mail addresses (M. Bonfanti), (V. Díaz-Zuccarini).

## Abstract

Patient-specific computational fluid-dynamics (CFD) can assist the clinical decision-making process for Type-B aortic dissection (AD) by providing detailed information on the complex intra-aortic haemodynamics. This study presents a new approach for the implementation of personalised CFD models using non-invasive, and oftentimes minimal, datasets *commonly* collected for AD monitoring. An innovative way to account for arterial compliance in rigid-wall simulations using a lumped capacitor is introduced, and a parameter estimation strategy for boundary conditions calibration is proposed. The approach was tested on three complex cases of AD, and the results were successfully compared against invasive blood pressure measurements. Haemodynamics results (e.g. intraluminal pressures, flow partition between the lumina, wall shear-stress based indices) provided information that could not be obtained using imaging alone, providing insight into the state of the disease. It was noted that small tears in the distal intimal flap induce disturbed flow in both lumina. Moreover, oscillatory pressures across the intimal flap were often observed in proximity to the tears in the abdominal region, which could indicate a risk of dynamic obstruction of the true lumen.

This study shows how combining commonly available clinical data with computational modelling can be a powerful tool to enhance clinical understanding of AD.

## 1 INTRODUCTION

Aortic dissection (AD) is a life-threatening vascular condition with high morbidity and mortality rates [1]. AD is characterized by the separation of the layers of the aortic wall: a tear in the intima layer allows the blood to flow within the aortic wall inducing the formation of two flow channels, the true (TL) and false lumen (FL), separated by an intimal flap (IF) [2].

Dissections not involving the ascending aorta are commonly managed with best medical treatment (BMT) in the absence of complications, such as end-organ malperfusion, rupture, refractory pain or hypertension. They include the ‘classic’ Type-B dissections (involving the descending thoracic aorta), the ‘Arch B’ dissections (involving the aortic arch and the descending thoracic aorta) [3] and the ‘residual post Type-A’ dissections (involving the aortic arch and the descending thoracic aorta after the surgical replacement of the ascending thoracic aorta). In this manuscript we conventionally report all three conditions as ‘Type-B ADs’. BMT alone is associated with poor long-term prognosis because up to 50% of the patients will develop complications requiring invasive management [4]. The identification of patients at risk of developing adverse events at an early stage would allow them to undertake elective endovascular treatment (TEVAR) in the acute or subacute phase, avoiding the challenges of chronic repair procedures [5] and the risks associated to emergency interventions [6]. Management and treatment of AD are highly patient-specific and morphological features, flow patterns, pressures, velocity and shear rates are extremely important features for this pathology. Hence, patient-specific computational fluid-dynamics (CFD) may lead to objective and quantifiable predictors of adverse outcomes and assist the clinical decision-making around the treatment of Type-B ADs by providing detailed information on the complex intra-aortic haemodynamics.

In a clinical scenario, the CFD modeller has to deal with the issue of incomplete and often noisy datasets to inform the computational models. In fact, practical, ethical and physical reasons prevent the acquisition of complete datasets necessary to construct fully “patient-specific” models [7], and adequate modelling assumptions have to be made to “fill the gaps”. For instance, a challenging and important task is the description of the boundary conditions (BCs). Since pressure and flow data are often unknown at the boundaries of the model, lumped parameters models (i.e. Windkessel models) coupled to the 3D domain are commonly employed to describe the pressure-flow relation at the boundaries due to the distal vasculature not included in the model. However, the calibration of the Windkessel models is not an easy exercise, and different strategies are proposed in the literature. Some strategies adopt only haemodynamic data taken from the literature [8], others make use of patient-specific flow waves integrated with literature-based pressure waves [7], while more advanced approaches use sophisticated techniques such as Kalman filters to assimilate boundary flow waves acquired with PC-MRI [9]. However, relying exclusively on literature data does not lead to the development of accurate personalised models and advanced clinical data, such as PC-MRI data, are not often acquired during routine monitoring.

In the case of ADs, the task of the modeller is further complicated by complex geometries and by the compliance of the arterial wall. As shown by Rudenick et al. [10] via a lumped parameter model and confirmed by Bonfanti et al. [11] with a CFD model, the wall compliance can indeed have a significant effect on the pressure in the FL, in particular when the size of the connecting tears is small. However, the majority of published AD models [12–16] assumes rigid-walls and neglects compliance effects on the predicted pressures. Advanced CFD models of AD account for the motion of the vessel walls using fluid-structure interaction (FSI) or moving boundary approaches [11,17–19] which, in order to be patient-specific, need to be informed by non-routine displacement data obtained, for example, via cine-MRI.

Thus, there is a need for cardiovascular CFD simulations to be further developed to account for the incomplete datasets commonly available for clinical translation. In this study, we present a computational framework for blood flow simulations of ADs which employs minimal datasets commonly acquired during routine monitoring. Lumped parameter models are used to simulate the aortic compliance and distal vasculature, and their parameters are calibrated with a new procedure which allows obtaining haemodynamic results in agreement with the available clinical data. The proposed framework is applied to 3 cases of complex Type-B ADs and the results are compared against invasive blood pressure measurements (IBPM), acquired in both the TL and FL, for validation purposes.

Haemodynamic results for the 3 cases are presented and analysed to enhance the clinical understanding of the disease. Finally, the applicability of the proposed framework and its limitations are discussed.

## 2 METHODS

### 2.1 Clinical dataset

The datasets of 3 patients (Table 1) with chronic AD was acquired as part of an ethically approved protocol at San Camillo-Forlanini Hospital (Rome, Italy) and included contrast-enhanced computed tomography (CT) scans, Doppler ultrasound measurements and IBPM.

**Table 1.** Details of the patients included in the study.

| Patient | Gender | Age | CO<br>[l/min] | HR<br>[bpm] | Inlet systolic/diastolic pressures<br>[mmHg] |
| --- | --- | --- | --- | --- | --- |
| #1 | M | 67 | 4.5 | 55 | 128/70 |
| #2 | M | 42 | 7.5 | 75 | 125/72 |
| #3 | F | 73 | 5.3 | 76 | 152/80 |

Doppler ultrasonography allowed the measurement of the cardiac output (CO) and heart rate (HR) of each patient as reported in Table 1. The IBPM allowed the measurement of the systolic and diastolic pressures at multiple intra-aortic locations both in the TL and FL.

Clinical CT scans covered the entire patients’ trunk from the supra-aortic vessels to the proximal iliac arteries. The resolution of the CT scans (max inter-slice distance = 1 mm, min in-plane resolution = 0.88 mm) was sufficiently high to visualise the IF and the main communicating tears between the TL and FL. The AD geometries were reconstructed with the image-processing software Simpleware ScanIP (Synopsys, USA).

Contrast-enhanced CT scans were analysed to detect evidence of malperfusion, as commonly done in clinical practice [20]. Asymmetric kidney enhancement indicating a reduced perfusion of the left kidney was observed in Patient 3. The percentage difference between the mean greyscale values in the two kidneys was evaluated using Simpleware Scan IP and found equal to −29% for the left kidney. No evidence of malperfusion was observed in the other patients.

### 2.2 Computational fluid dynamics

#### 2.2.1. Flow model and boundary conditions

The Navier–Stokes (NS) and continuity equations for 3D time-dependent flows were solved with the finite-volume-based CFD solver ANSYS-CFX 18.0 (ANSYS Inc., PA, USA). Blood was modelled as incompressible with a density of 1056 kg m^−3^ and non-Newtonian viscosity described by the Carreau–Yasuda model with parameters taken from Gijsen et al. [21]. The blood flow was considered laminar; based on the inlet area of the reconstructed aortae, the obtained mean Reynolds numbers varied from 665 to 1506 and the Womersley numbers from 20 to 32. The peak Reynolds numbers varied between 1972 and 2933 and were in all cases lower than the critical Reynolds numbers for transition to turbulence calculated following Peacock et al. [22] considering a viscosity of 4 × 10^−3^ Pa s.

A representative CFD model and its BCs is shown in Fig. 1a for Patient 1.

**Figure 1.**
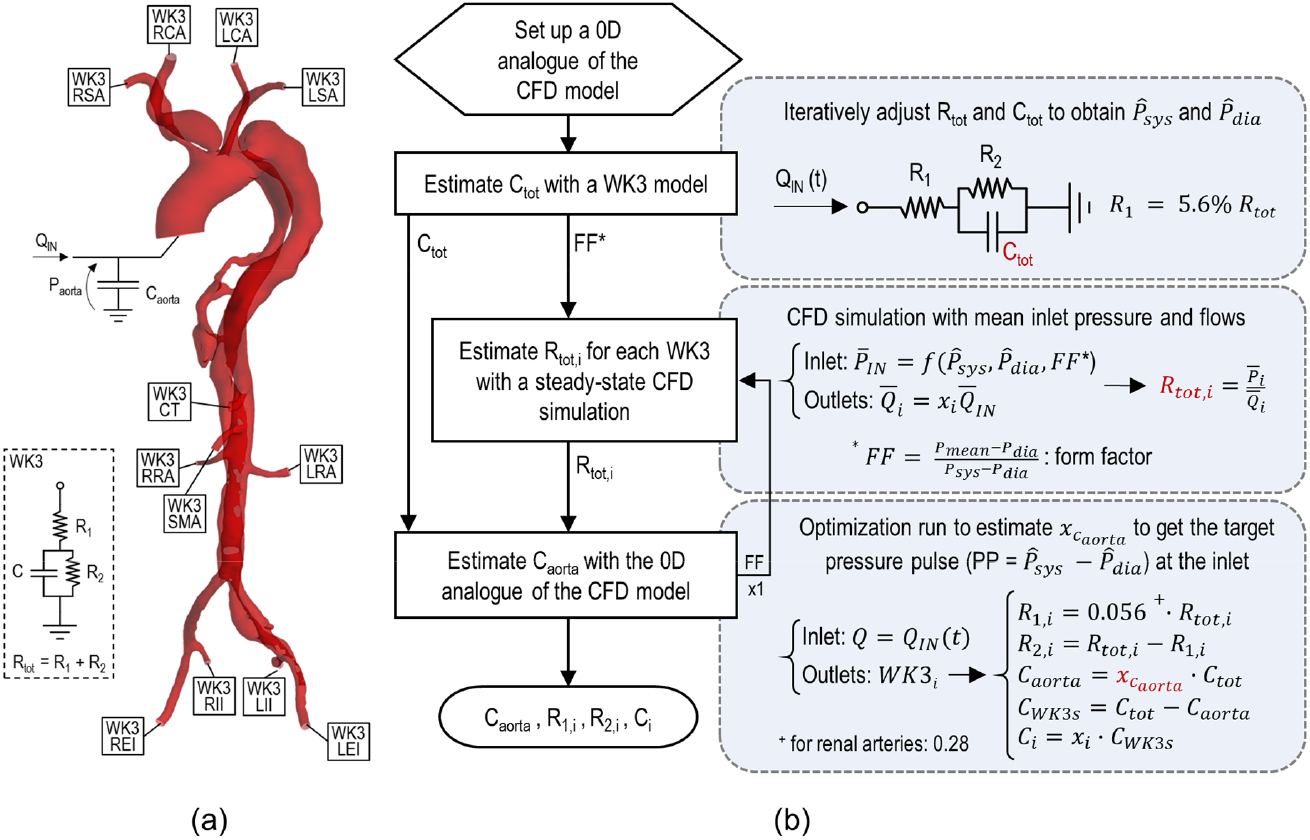
(a) Schematic of the CFD model and its boundary conditions. Patient 1 is shown as an example, (b) Flowchart of the procedure adopted for parameter calibration.

The fluid-structure interaction effects due to the compliance of the aorta were modelled in a lumped manner by introducing a capacitor (C_aorta_) before the inlet of the 3D model, following an approach similar to the one in Pant et al. [9].

As a result, the flowrate Q_3D_ prescribed at the inlet boundary of the 3D model is related to the inlet flow Q_IN_ via Eq. 1:

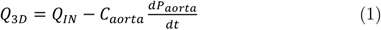

where P_aorta_ is the average pressure in the aorta. C_aorta_ is estimated as described in Section 2.2.2. Q_3D_ was prescribed based on a uniform velocity profile.

The inlet flowrate Q_IN_ was obtained by adjusting a typical ascending aorta blood flow waveform [23] to the patient-specific haemodynamic data acquired by Doppler ultrasound (i.e. CO, HR and systolic-to-diastolic duration ratio).

A three-element Windkessel model (WK3) was coupled to each outlet *i*. Hence, the outlet flow (Q_i_) and mean pressure (P_i_) are related by

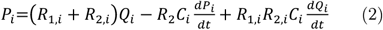

where R_1,i_ and R_2,i_ represent the proximal and distal resistances, and C_i_ is the compliance of the vasculature distal to outlet *i*. WK3 parameters were calibrated as detailed in Section 2.2.2.

The 3D model walls were assumed to be rigid with a no-slip condition.

#### 2.2.2 Parameter calibration procedure for model personalisation

The WK3 model parameters (i.e. R_1,i_ R_2,i_ and C_i_) and C_aorta_ were adapted to the specific patient following the procedure described below. The objectives of the calibration procedure were (i) to achieve physiological flow distribution at each outlet, and (ii) to obtain the *measured* systolic 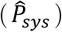 and diastolic 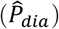 pressures at the inlet. In this study 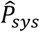 and 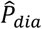 were available from IBPM, but they can also be estimated from non-invasive brachial pressure measurements as described, e.g., by Saouti et al. [24].

The fraction of the CO leaving each outlet (*x_i_*) was determined based both on available patient data and on the literature as follows: 30% of the CO was directed to the supra-aortic branches, as typically reported for AD patients [11,13], and distributed among them proportionally to the vessel cross-sectional area [25]. The mean flow exiting the celiac trunk (CT), the superior mesenteric artery (SMA) and the renal arteries (LRA, RRA) was determined according to Nakamura et al. [26] and Hall and Guyton [27].

The remaining mean flow was equally split between the common iliac arteries, 70% of which was directed to the external iliac arteries (REI, LEI) and the remaining 30% to the internal ones (RII, LII). In case of evidence of non-physiological CO distribution, the estimated values were adjusted accordingly. For example, the mean flow in the LRA of Patient 2 was reduced by 29% according to the available CT scans.

Fig. 1b illustrates the workflow adopted for the model calibration comprising the following steps:

1. The first step involves setting up a reduced-order analogue of the CFD model. The 3D aorta was divided in segments modelled as 0D-building-blocks made by *inertances* (*Ls*) and *resistances* (*Rs*). Ls were estimated from the geometry of the segments (i.e. mean cross-sectional area and length) [28], while the Rs were calculated as the ratio of the pressure drop to the flowrate at each segment as determined by a steady-state CFD simulation of the 3D AD model.
2. A WK3 analogue of the entire vascular system was used to estimate the total arterial compliance C_tot_, following the method described by Les et al. [29]. WK3 parameters C_tot_, R_1_ and R_2_ (R_1_/(R_1_ + R_2_) = 5.6% [29]) were iteratively varied until the pressure waveform P(t) obtained was bounded by the target 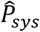 and 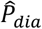, using Q_IN_(t) as input. An estimate of the form factor FF = (P_mean_-P_dia_)/(P_sys_-P_dia_) [30], necessary for Step 3, was calculated, where P_mean_, P_sys_ and P_dia_ are the mean, maximum and minimum values of P(t), respectively.
3. The total resistance R_tot,i_ = R_1,i_ + R_2,i_ of each WK3 was estimated via a steady-state CFD simulation of the AD model with the following BCs: the mean pressure 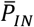 calculated from 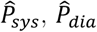 and FF was prescribed at the inlet, while the target mean flow (i.e. 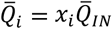) was imposed at the outlets as outflow condition. R_tot_,_i_ was calculated as the ratio of the predicted outlet pressure 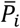 to 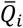.
4. The reduced-order model set up at Step 1 was used at this stage as an analogue of the CFD model to determine C_aorta_ as a fraction *x_c_aorta__* of the estimated total compliance (*C_aorta_* = *x_c_aorta__C_tot_*). The compliance attributed to the outlets (*C_WK3s_*) was calculated as the difference between C_tot_ and C_aorta_ and distributed to each WK3 proportionally to the mean flow 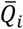. Values of R_tot,i_ from Step 3 were used for the WK3s, assuming a R_1,i_/R_tot,i_ ratio equal to 28% for the renal arteries and 5.6% for the other outlets, respectively, following Les et al. [29]. The fraction *x_c_arota__* was estimated using an optimization algorithm (Broyden–Fletcher–Goldfarb– Shanno algorithm) in order to obtain the target pulse pressure 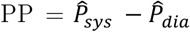 at the inlet of the reduced-order model. The governing equations were solved with a back-ward-differentiation scheme using the software 20-sim (Controllab Products B.V., Enschede, The Netherlands).
5. Finally, Steps 3 and 4 were reiterated using the FF calculated from the inlet pressure waveform obtained at Step 4. The obtained model parameters C_aorta_, R_1,i_ R_2,i_ and C_i_ were used for the final CFD model.

#### 2.2.3 Numerical simulations and postprocessing

The NS equations were spatially and temporally discretised with a high resolution advection scheme [31] and a second-order backward Euler scheme, respectively, using a uniform time-step of 1 ms, which was deemed sufficient for time-step size independent results. Time-derivative terms in Eqs. 1 and 2 were discretized with a first-order backward Euler approach.

Computations were run in parallel mode (approximately 35,000 nodes per partition) on the high-performance computing cluster of the Department of Computer Science, at University College London. Average computational time per cardiac cycle was about 15 hours. Simulations were run until reaching a periodic steady-state which took 3 to 4 cardiac cycles, and the last cycle was used for the analysis of results.

Post-processing was performed using CFD-Post (ANSYS Inc.) and MATLAB (Mathworks, MA, USA). Time-averaged haemodynamic indices, such as Time-Averaged Wall Shear Stress (TAWSS), Oscillatory Shear Index (OSI) and Relative Residence Time (RRT), were calculated according to Gallo et al. [32]. TAWSS is the average of the viscous tangential stresses on the vessel wall over a cardiac cycle; OSI indicates regions were the instantaneous wall shear stress deviates from the main flow direction over large parts of the cardiac cycle and is an index of disturbed flow [32]; RRT indicates regions of blood stagnation and can be used as a proxy of particle deposition [8].

The pressure difference between the TL and FL (△P = P_TL_ - P_FL_) was evaluated over several crosssectional planes along the dissection, as shown in Figures 4–6, at three different instants of the cardiac cycle (i.e. mid-acceleration, peak systole, mid-deceleration). A positive ΔP indicates higher pressure in the TL, whereas negative values mean higher pressures in the FL.

#### 2.2.4 Mesh

The AD geometries were discretised with ICEM-CFD (ANSYS Inc.) using a tetrahedral mesh in the core region and seven prism layers at the wall boundaries, as previously done for similar aortic geometries [11,17].

A mesh independence study was carried out to select the parameters to be used for grid creation (e.g. element size, curvature/proximity refinement settings). Coarse, medium and fine meshes were created for Patient 3 corresponding to approximately 750 K, 1.7 M and 3 M elements, respectively. Pressure and flow waves at the outlets were compared resulting in a maximum difference of 2.4% and 1.0%, respectively, when comparing the coarse to the medium mesh, and 1.4% and 1.0% when comparing the medium to the fine grid. The TAWSS, OSI and RRT maps obtained with the three grids were qualitatively similar, with an average difference of only 0.032 Pa, 0.021 and 2.11 Pa^−1^, respectively, when comparing the medium to the fine grid, and of 0.20 Pa, 0.025 and 2.41 Pa^−1^ when comparing the coarse to the medium grid. Following this analysis, the medium grid was deemed sufficient for simulation purposes, and the same parameters were used for the discretization of Patient 1 and 2 geometries. The number of elements of the generated grids varied between 1.5 and 2.4 M, depending on the size and structure of the models.

Further simulations were run for Patients 1 and 2 on coarser grids with approximately half number of elements, resulting in a maximum difference of less than 5.1% and 2.3% when comparing the outlets’ flow and pressure waves, respectively, to the medium mesh results. The obtained TAWSS and OSI maps were qualitatively similar, with a maximum average difference of 0.35 Pa and 0.031 for the TAWSS and OSI, respectively. These differences are comparable with those obtained for Patient 3 when moving from a coarse to a medium grid, and prove the suitability of the selected parameters for the discretisation of the 3D geometries.

## 3 RESULTS

### 3.1 Aortic dissections anatomy

#### Patient 1

Patient 1 presents a Type-B AD originating from an entry tear (area ≅ 150 mm^2^) just distal to the LSA and extending to LEI (Fig. 2). A dissection flap can be noted in the brachiocephalic trunk (BT). Multiple tears are present along the dissection as indicated in Fig. 2. All the aortic branches originate from the TL. Lastly, an additional FL is observed at the level of the distal thoracic aorta (dashed green arrows in Fig. 2).

**Figure 2.**
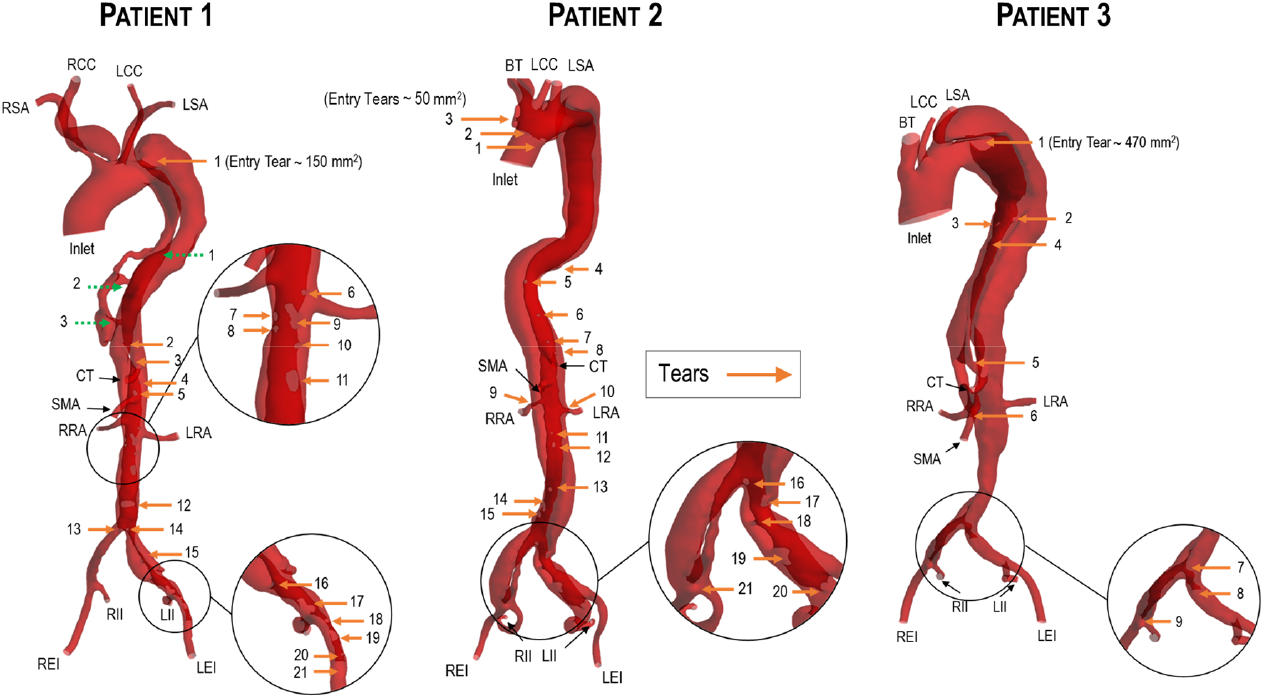
Geometry of the ADs as extracted from the CT scans for the three patients. Arrows indicate the location of the tears in the intimal flap.

#### Patient 2

Patient 2 presents a residual Type-B dissection following a surgical repair of a Type-A AD which required the replacement of the ascending aorta with a vascular graft. The residual dissection extends to the distal common iliac arteries. Three small entry tears are observed in the proximal part of the dissection (area of the tears ≅ 50 mm^2^) and further multiple connecting tears are present in the IF as illustrated in Fig. 2. The dissection involves BT, RRA and LRA which are perfused by both the TL and the FL, while the other aortic branches originate from the TL.

#### Patient 3

Patient 3 has a Type-B AD which extends from just distal to the LSA to the common iliac arteries. A large entry tear of about 470 mm^2^ is present in the proximal part of the dissection, about 45 mm distal to the LSA. The RRA and SMA originate from the TL, whereas the LRA from the FL. A dissection flap is observed in the CT which is perfused by both the TL and the FL. Multiple tears are present in the IF as illustrated in Fig. 2 and, as in most cases, they are in proximity to the origin of intercostal, visceral or renal arteries from the FL.

### 3.2 Personalised boundary conditions

The model parameters estimated using the proposed calibration procedure are listed in Tab. 2. The calibration objectives were met closely as reported in Tab. 3, with a maximum difference between the obtained and target values of 0.5% for the flow split and 3 mmHg for the inlet pressures.

**Table 2.** Comparison between target and computed values for flow split and inlet pressures for the 3 patients.

| Pat. # | | $x_{RSA}$ | $x_{LSA}$ | $x_{RCC}$ | $x_{LCC}$ | $x_{BT}$ | $x_{CT}$ | $x_{SMA}$ | $x_{RRA}$ | $x_{LRA}$ | $x_{REI}$ | $x_{LEI}$ | $x_{RII}$ | $x_{LII}$ | $P_{sys}$ | $P_{dia}$ |
| --- | --- | --- | --- | --- | --- | --- | --- | --- | --- | --- | --- | --- | --- | --- | --- | --- |
| 1 | CFD | 8.1 | 6.0 | 10.1 | 6.1 | n/a | 19.5 | 14.8 | 10.4 | 10.6 | 5.1 | 5.1 | 2.0 | 2.1 | 130.8 | 69.7 |
|  | Target | 8.0 | 6.0 | 10.0 | 6.0 | n/a | 20.0 | 15.0 | 10.5 | 10.5 | 5.0 | 5.0 | 2.0 | 2.0 | 128.0 | 70.0 |
|  | Diff. | 0.1 | 0.0 | 0.1 | 0.1 | n/a | -0.5 | -0.2 | -0.1 | 0.1 | 0.1 | 0.1 | 0.0 | 0.1 | 2.8 | -1.3 |
| 2 | CFD | n/a | 7.8 | n/a | 3.7 | 18.5 | 13.4 | 10.5 | 10.5 | 10.5 | 8.8 | 8.8 | 3.7 | 3.8 | 125.6 | 69.0 |
|  | Target | n/a | 7.9 | n/a | 3.8 | 18.3 | 13.5 | 10.4 | 10.5 | 10.5 | 8.8 | 8.8 | 3.8 | 3.8 | 125 | 72.0 |
|  | Diff. | n/a | -0.1 | n/a | -0.1 | 0.2 | -0.1 | 0.1 | 0.0 | 0.0 | 0.0 | 0.0 | -0.1 | 0.0 | 0.6 | -3.0 |
| 3 | CFD | n/a | 7.1 | n/a | 4.1 | 19.0 | 16.2 | 13.5 | 10.5 | 7.5 | 7.7 | 7.8 | 3.3 | 3.3 | 152.3 | 81.7 |
|  | Target | n/a | 7.1 | n/a | 4.1 | 18.8 | 16.4 | 13.6 | 10.5 | 7.5 | 7.7 | 7.7 | 3.3 | 3.3 | 152.0 | 80.0 |
|  | Diff. | n/a | 0.0 | n/a | 0.0 | 0.2 | -0.2 | -0.1 | 0.0 | 0.0 | 0.0 | 0.1 | 0.0 | 0.0 | 0.34 | 1.7 |

\* $x_i$ is the mean flow split in %, $P_{sys}$ and $P_{dia}$ are the inlet systolic and diastolic pressures in mmHg.

**Table 3.**
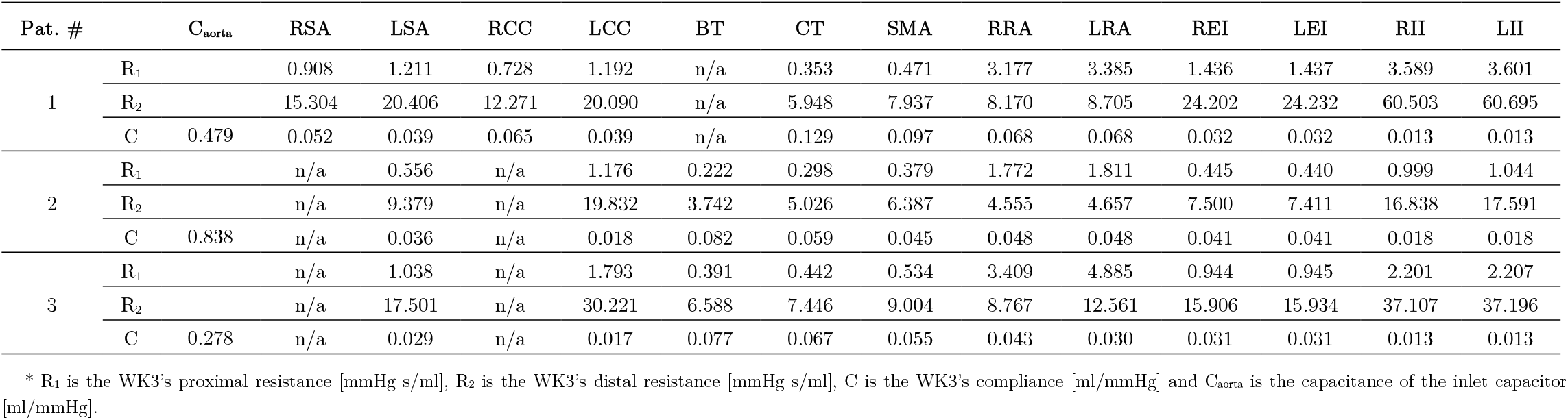
Parameters obtained using the proposed calibration procedure for patient specific boundary conditions.

The inlet capacitors accounted for 42.6, 64.9 and 40.6% of C_tot_ estimated for the 3 models, respectively, consistent with the findings reported by Ioannou et al. [33] according to which approximately 50% for the total arterial compliance is located in the proximal thoracic aorta [34].

The models allowed physiological inlet pressure waveforms to be obtained with FF values in the range of 0.49-0.51, as expected at the aortic root level [35].

### 3.3 Comparison against invasive blood pressure measurements

A comparison between the systolic and diastolic pressures obtained with the CFD models and those recorded via IBPM at various intra-aortic locations, both in the TL and FL, is shown in Fig. 3. The two sets of data compare well, with a mean difference (± standard deviation) between the computational and clinical data of 4 ± 2 mmHg for Patient 1, 3 ± 1 mmHg for Patient 2 and 4 ± 3 mmHg for Patient 3. These differences are in accordance with the accuracy expected for IBPM, as reported by Romagnoli et al. [36], and give confidence on the reliability of the simulation results. It should be noted that while the pressure field calculated by the simulation represents a heartbeat as determined by the applied BCs, the clinical measurements are inevitably affected by cycle-to-cycle variability; this represents a source of error which should be considered when comparing clinical and simulation data.

**Figure 3.**
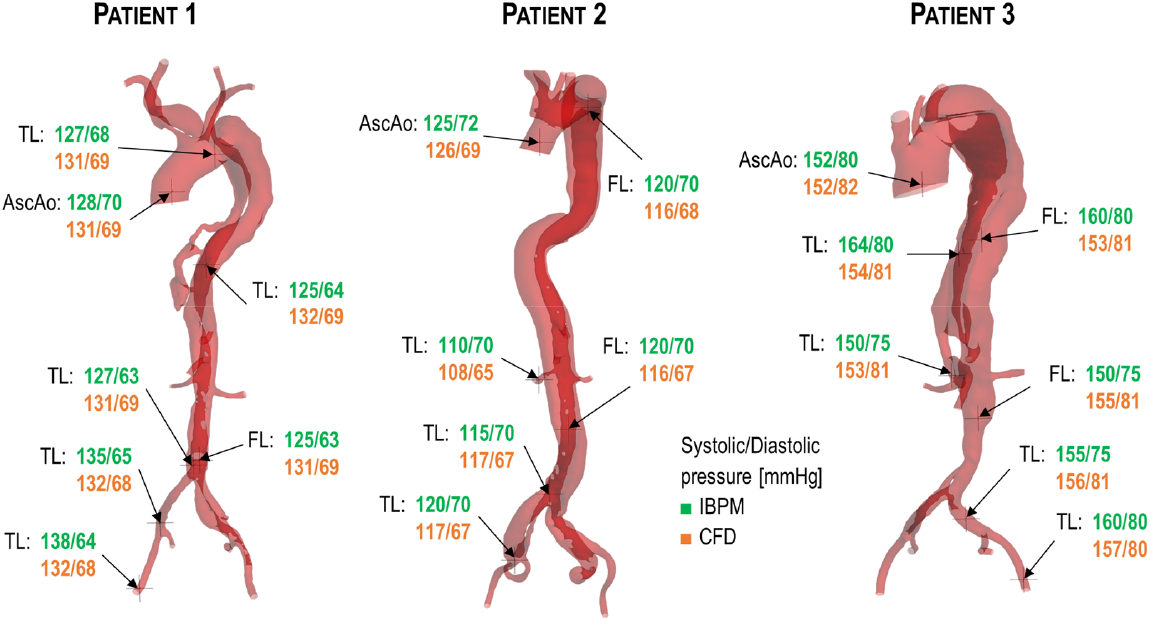
Comparison between computational (orange) and invasive intra-aortic blood pressure measurements (IBPM) (green).

### 3.4 Blood flow dynamics

#### Patient 1

Computational haemodynamic results obtained for Patient 1 are presented in Fig. 4. The computed pressure field at peak systole (Fig. 4a) shows a pressure gradient along the length of the TL of 15 mmHg (from the entry tear to the main exit tear at the iliac bifurcation) which is larger than the values reported for ‘undissected’ aortae (approx. 5 mmHg [12]). This is due to the higher hydraulic resistance of the TL due to its reduced cross-sectional area, which ultimately leads to a higher hydraulic load placed on the left ventricle. Only a small proportion of the total aortic flow enters the FL via the main entry tear (15%), while most of the blood flows through the TL perfusing the main aortic branches. Pressures in the TL are higher than in the FL in the proximal part of the dissection (Fig. 4b), with a maximum ΔP at peak systole of 10 mmHg. The pressure difference ΔP between the TL and FL decreases along the dissection and negative values can be noted in the distal region, just above the CT (at the mid-deceleration systolic phase), and above RRA, LRA and SMA (at peak systole). Such ΔP variations may lead to dynamic compression of the TL due to IF motion, and consequently hinder the blood flow in these important aortic branches originating from the TL.

**Figure 4.**
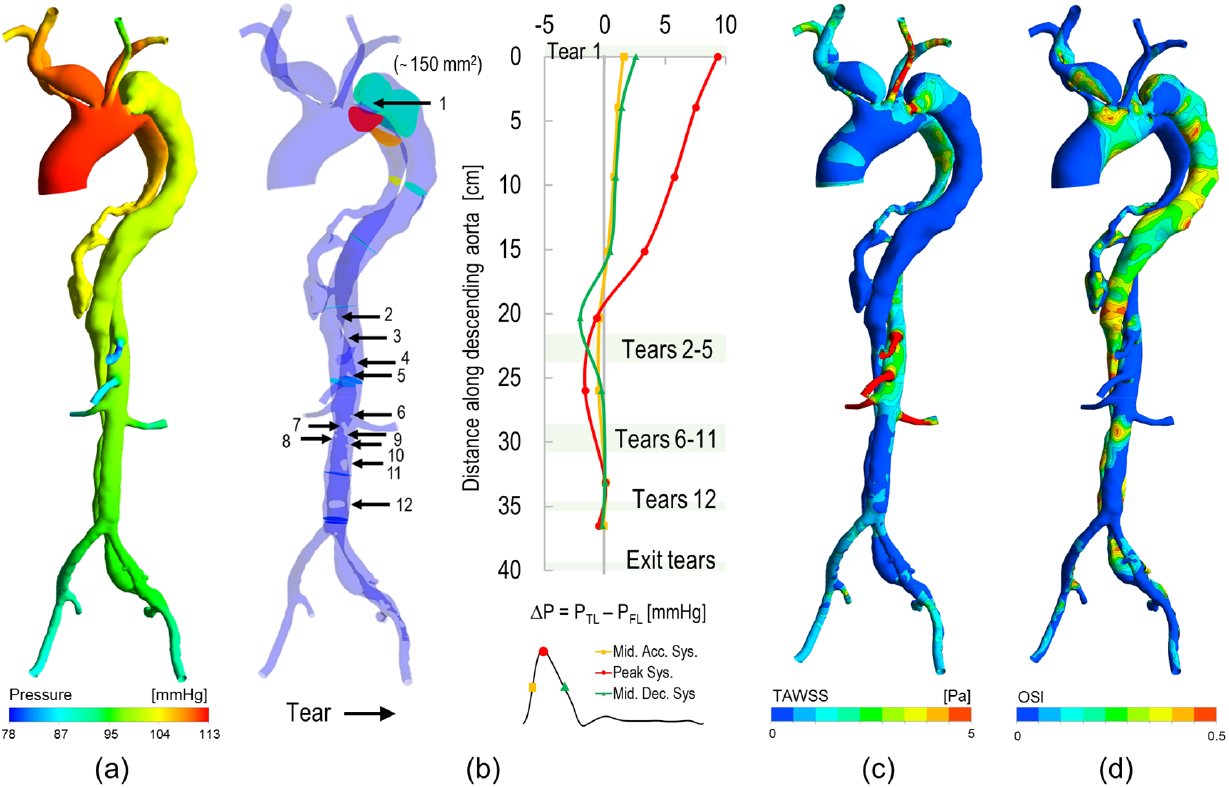
Haemodynamic results for Patient 1. (a) Pressure field obtained at peak systole; (b) pressure difference (△P) between the TL and the FL calculated at three different phases of the cardiac cycle; (c) Time-averaged wall shear stress (TAWSS) distribution; (d) Oscillatory shear index (OSI) distribution.

TAWSSs in the TL (range 0.5-12 Pa, Fig. 4c) are higher than those reported for healthy aortae (<2 Pa [37]); focal regions of increased TAWSS are observed in the TL where the cross-sectional area is smaller and at the location of intimal flap tears (e.g. entry tear TAWSS = 7.5 Pa) which could represent a risk of tear expansion. On the contrary, lower TAWSSs are observed in the FL (< 2 Pa) due to lower velocities and larger crosssectional areas. Here, slow and disturbed flow leads to high values of OSI, as can be seen in Fig. 4d. High OSI values are also observed on the TL wall next to IF tears.

#### Patient 2

Computational haemodynamic results obtained for Patient 2 are presented in Fig. 5. A large pressure gradient along the TL can be noted at peak systole, leading to a pressure drop of 21 mmHg from the proximal TL to the iliac bifurcation. This is due to the very small crosssectional area of the TL, which is almost completely compressed by the FL in the distal part of the thoracic aorta. Even if only small entry tears connect the TL to the FL, the blood flow splits almost equally between the TL and FL (44% in the FL, 56% in the TL) due to the high hydraulic resistance of the TL. The small entry tears lead to higher proximal pressures in TL than in the FL throughout the systolic phase, with a maximum ΔP of 15 mmHg at peak systole. The ΔP decreases in the distal descending aorta, becoming negative at the level of the abdominal branches through half of the systolic phase, and hence presenting a risk of dynamic obstruction of the already narrow TL. Nonetheless, perfusion of the kidneys does not appear to be at risk since a significant portion of the renal blood flow comes from the FL (i.e. 73 and 48% for LRA and RRA, respectively).

**Figure 5.**
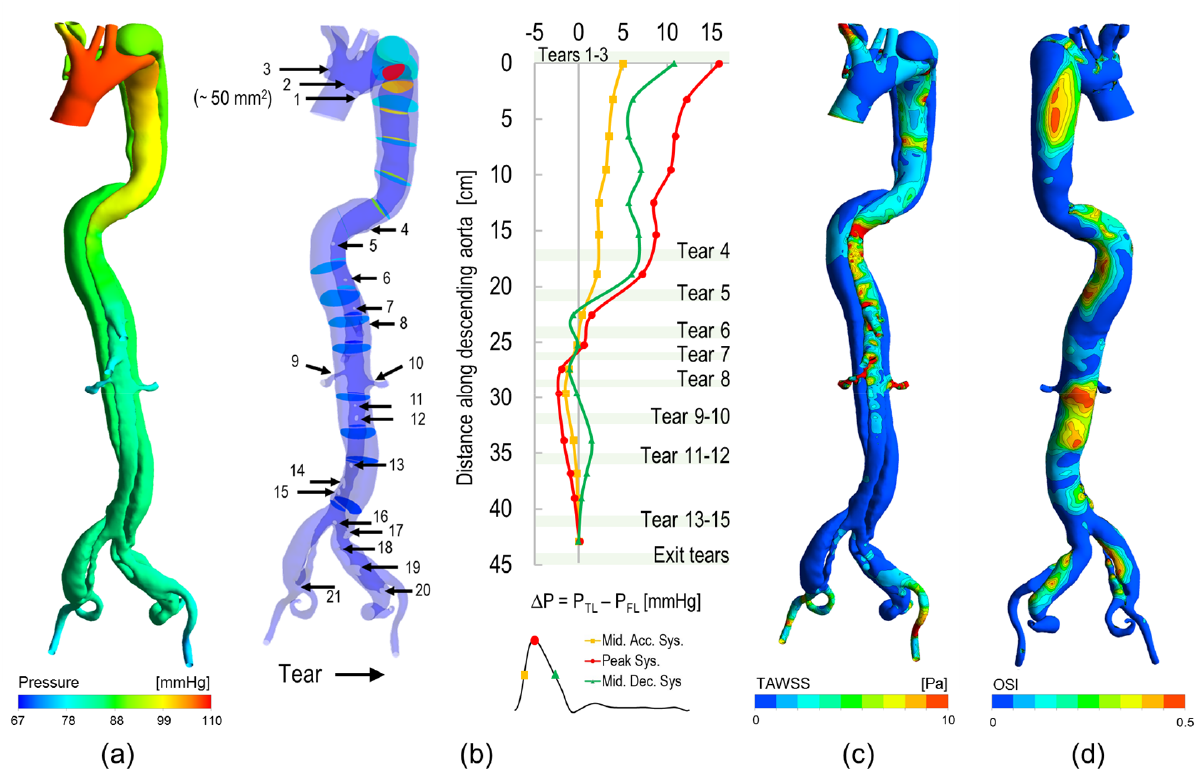
Haemodynamic results for Patient 2. (a) Pressure field obtained at peak systole; (b) pressure difference (△P) between the TL and the FL calculated at three different phases of the cardiac cycle; (c) Time-averaged wall shear stress (TAWSS) distribution; (d) Oscillatory shear index (OSI) distribution.

Extremely high TAWSS values are obtained at the entry tears (up to 38 Pa) and where the TL cross-section is minimum (24 Pa). The marked bending of the aorta in two points leads to regions of flow separation and reattachment characterised by high values of OSI as can be seen on the FL wall. The presence of tears induces disturbed flow in both the TL and FL as indicated by elevated OSI values in the distal abdominal aorta.

#### Patient 3

Haemodynamic results for Patient 3 are presented in Fig. 6. The dissection is characterized by a large entry tear which diverts most of the aortic flow (88%) towards the FL. The pressure gradient along the aorta at peak systole is modest (about 7 mmHg) due to the large cross-sectional area of the FL and the little flow going through the smaller TL.

**Figure 6.**
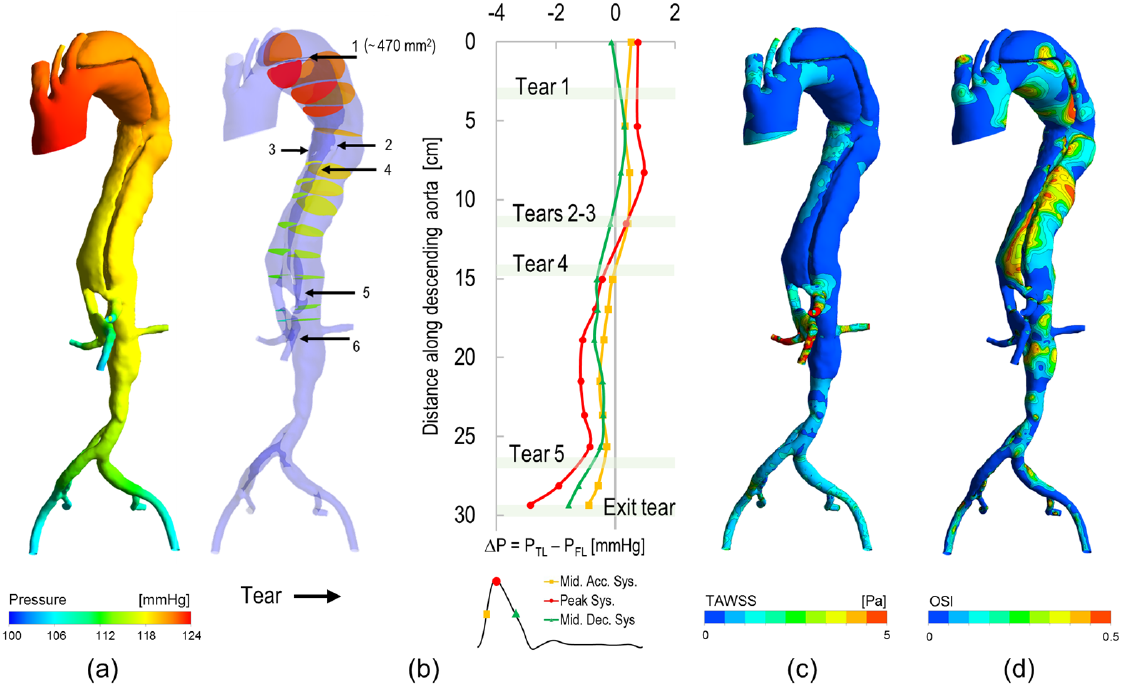
Haemodynamic results for Patient 3. (a) Pressure field obtained at peak systole; (b) pressure difference (△P) between the TL and the FL calculated at three different phases of the cardiac cycle; (c) Time-averaged wall shear stress (TAWSS) distribution; (d) Oscillatory shear index(OSI) distribution.

Due to the large entry tear, the pressure is almost equal in the TL and FL starting from the proximal part of the dissection (Fig. 6b). The pressure difference ΔP between the TL and FL becomes negative in the region of the abdominal branches where the constricted TL results in higher blood velocities and lower pressures (i.e. Venturi effect). However, because of the thick IF in this region, the risk of compression of the TL due to IF motion is low.

The TAWSSs acting on the aortic wall are physiological (< 2 Pa) while regions of high OSI can be noted in both the TL and FL, mainly due to the irregular surface of the vessel. In Fig. 7b, a large FL partial thrombosis compressing the TL can be noted in the abdominal region just above important branches originating from the TL (i.e. SMA, RRA). In the same region, simulation results show high values of RRT on the TL wall (Fig. 7a), which have been found to correlate positively with thrombus formation [8]; further thrombus growth could hinder the flow in the TL and cause a risk of end-organ malperfu-sion.

**Figure 7.**
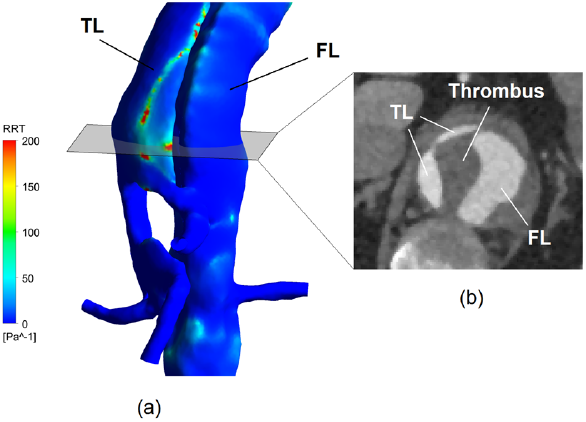
(a) Relative Residence Time (RRT) in the abdominalaorta ofPatient 3. High values of RRT can be noted on the wall of the TL. RRT is proportional to the residence time of blood particle near the wall, and it has been correlated to regions of thrombosis, (b) CT scan showing a large FL partial thrombosis compressing the TL.

## 4 DISCUSSION

One of the main challenges engineers face in translating patient-specific modelling into the clinic is the ability to reproduce the specific patient condition accurately enough to be clinically meaningful using data that are acquired routinely and non-invasively. In this work, we presented a computational framework that allows the implementation of blood flow simulations of AD informed by datasets commonly acquired during AD monitoring. The framework was tested on three complex cases of dissection of the descending thoracic aorta (and the aortic arch, in two cases), and the results were compared against invasive blood pressure measurements (IBPM).

### 4.1 Modelling approach: boundary conditions

The proposed computational framework features dynamic BCs which enable obtaining physiological pressure and flow waves in the CFD domain. State-of-the-art Windkessel models were used as outflow BCs in order to represent the distal vasculature, which have been shown to be superior to other BCs strategies by Pirola et al. [37], while at the inlet a novel approach was adopted to represent the compliance of the aorta via a lumped parameter.

Pressure is an important variable that can be extracted from CFD simulations and it is particularly relevant for AD, where high pressures in the FL could indicate a risk of expansion and rupture. Such pressures cannot be measured non-invasively. However, without accounting for aortic wall compliance, it is not possible to obtain physiologically reasonable results in terms of pressure values and therefore match measured invasive data, if available. On the other hand, moving wall simulations that inherently account for vessel compliance with FSI or moving boundary techniques pose additional difficulties. Besides being computationally expensive and challenging in case of large deformations in complex AD geometries, these models need to be informed by imaging data of the vessel wall position, which are not routinely acquired.

The lumped capacitor adopted as inlet BC in this study allows to account for aortic compliance avoiding moving wall simulations. This approach is similar to the one proposed by Pant et al. [9], and it is based on the hypothesis that, since the majority of the arterial compliance is located in the proximal thoracic aorta [33], “lumping” it at the inlet would not significantly affect the haemodynamics in the descending thoracic aorta, which is the focus of Type-B AD simulations. In the absence of complete datasets, including wall deformation data, and in order to avoid moving wall simulations, this method represents the best compromise for obtaining physiological pressures in patient-specific simulations. The model results were compared against *in vivo* IBPM, and the good match achieved validated the approach.

### 4.2 Calibration strategy

The proposed strategy for the calibration of the BCs was tested on all 3 AD cases. Target values for pressures and flow were matched reasonably well. The calibration objectives (i.e. flow distribution and inlet systolic/diastolic pressure) are the same as those commonly adopted when tuning Windkessel parameters for blood flow simulations [13,29,38]. However, calibration strategies previously reported in the literature often rely on iterative procedures in the context of 1D models [38,39] which would be computationally prohibitive if directly applied to 3D models. With the proposed method, by combining time-dependent simulations on a reduced-order model and steady-state simulations on the actual 3D model, we were able to reduce the computational time needed for the calibration and, at the same time, account for the haemodynamic resistance of the pathological aortae which likely affects the flow distribution.

### 4.3 Haemodynamic results

Simulations provided information that could not be obtained by imaging alone, such as the dynamics of the intra-luminal pressure, TAWSS, OSI and RRT distributions, that can be used to predict the progression of AD as discussed in the results section. As a general observation, it was noted that tears in the distal IF induce disturbed flow both in the TL and FL, as can be seen from high OSI values. Moreover, oscillatory transmural pressures across the IF were often observed in the proximity of tears in the abdominal region, which could lead to dynamic obstruction of the TL. Furthermore, it was noted that flow partition between the TL and the FL does not depend only on the size of the main entry tears, but also on the shape of the TL. For example, in patient 2, even if the entry tears are small, the blood flow splits almost equally between the two lumina due to the small cross-sectional area of the TL.

### 4.4 Limitations and applicability

The proposed approach presents some limitations deriving from the approximation of the vessel compliance via a lumped capacitor at the inlet and the assumption of rigid walls.

Firstly, by moving the compliance from the proximal aorta to the inlet, accurate blood flow predictions in the descending aorta can be made. This will be effective for the study of Type-B dissections, but may not be suitable for dissections involving the ascending aorta.

Secondly, studies based on lumped parameter models investigating the effect of wall elasticity on AD haemodynamics [10] showed that, in case of small communications between the two lumina, the effect of wall elasticity on pressure predictions can be relevant. In particular, it was observed that the FL pressure wave is slower than the TL one in the case of small tears and distensible walls, leading to a time-shift between the TL and FL waveforms that can significantly impact the instantaneous transmural pressure across the IF. This effect was confirmed with a 3D compliant model of an AD lacking re-entry tears [11]. On the other hand, the time-shift between the pressure waves tends to zero in the presence of at least one large tear [10,40], thus in this case a rigid-wall model would be good enough for assessing the pressure difference between the lumina [41].

Lastly, rigid-wall models do not allow an accurate description of the local effects of the motion of IF on the flow field [17] and thus may not be suitable in acute settings where the IF is highly mobile.

In summary, the proposed approach is applicable for the study of ADs where the region of interest is the descending aorta, with large or multiple communications between the TL and the FL, and with a fairly rigid intimal flap.

## 5 CONCLUSIONS

This paper presents a new method for the implementation of personalised CFD models using non-invasive and minimal datasets routinely acquired for AD monitoring, bringing CFD simulations closer to the clinic. The approach was tested on three complex cases of Type-B AD, and the results were positively compared against IBPM, specifically acquired in this study for validation purposes only, but not necessary for the implementation of the models. An innovative way to account for wall compliance via a lumped parameter was investigated, and a personalisation strategy for selecting the model parameters was proposed.

This study shows how combining commonly available clinical data with computational modelling can be a powerful tool to increase clinical understanding of aortic dissection.

## ETHICAL APPROVAL

The study was ethically approved (A.O. San Camillo Forlanini, Prot. n. 900/CE Lazio 1). The patients signed the appropriate consent form.

## ACKNOWLEDGEMENTS

This project was supported by the European Union’s Horizon 2020 research and innovation programme (Marie Sklodowska-Curie GA No. 642612, www.vph-case.eu); the Well-come/EPSRC Centre for Interventional and Surgical Sciences (WEISS) (203145Z/16/Z); and the Leverhulme Trust (Senior Research Fellowship No. RF-2015-482).

The authors would also like to thank the Department of Computer Science of University College London for the high-performance computing cluster resources used to perform the simulations.

## REFERENCES

[1] Nienaber CA, Clough RE, Sakalihasan N, Suzuki T, Gibbs R, Mussa F, et al. Aortic dissection. Nat Rev Dis Prim 2016;2:16053. doi:10.1038/nrdp.2016.53.

[2] Strayer RJ, Shearer PL, Hermann LK. Screening, evaluation, and early management of acute aortic dissection in the ED. Curr Cardiol Rev 2012;8:152–7.

[3] Trimarchi S, de Beaufort HWL, Tolenaar JL, Bavaria JE, Desai ND, Di Eusanio M, et al. Acute aortic dissections with entry tear in the arch: A report from the International Registry of Acute Aortic Dissection. J Thorac Cardiovasc Surg 2019;157:66–73. doi:10.1016/J.JTCVS.2018.07.101.

[4] Kamman A V, Brunkwall J, Verhoeven EL, Heijmen RH, Trimarchi S. Predictors of aortic growth in uncomplicated type B aortic dissection from the Acute Dissection Stent Grafting or Best Medical Treatment (ADSORB) database. J Vasc Surg 2017;65:964–71. doi:10.1016/j.jvs.2016.09.033.

[5] Erbel R, Aboyans V, Boileau C, Bossone E, Di Bartolomeo R. 2014 ESC Guidelines on the diagnosis and treatment of aortic diseases. Eur Heart J 2014;35:2873–2926. doi:10.1093/eurheartj/ehu281.

[6] Nienaber CA, Rousseau H, Eggebrecht H, Kische S, Fattori R, Rehders TC, et al. Randomized Comparison of Strategies for Type B Aortic Dissection. Circulation 2009;120.

[7] Romarowski RM, Lefieux A, Morganti S, Veneziani A, Auricchio F. Patient-specific CFD modelling in the thoracic aorta with PC-MRI-based boundary conditions: A least-square three-element Windkessel approach. Int j Numer Method Biomed Eng 2018;34:e3134. doi:10.1002/cnm.3134.

[8] Xu H, Piccinelli M, Leshnower BG, Taylor WR, Veneziani A. Coupled Morphological–Hemodynamic Computational Analysis of Type B Aortic Dissection: A Longitudinal Study. Ann Biomed Eng 2018. doi:10.1007/s10439-018-2012-z.

[9] Pant S, Fabrèges B, Gerbeau J-F, Vignon-Clementel IE. A methodological paradigm for patient-specific multi-scale CFD simulations: from clinical measurements to parameter estimates for individual analysis. Int j Numer Method Biomed Eng 2014;30:1614–48. doi:10.1002/cnm.2692.

[10] Rudenick PA, Bijnens BH, Segers P, García-Dorado D, Evangelista A. Assessment of wall elasticity variations on intraluminal haemodynamics in descending aortic dissections using a lumped-parameter model. PLoS One 2015;10:1–17. doi:10.1371/journal.pone.0124011.

[11] Bonfanti M, Balabani S, Greenwood JP, Puppala S, Homer-Vanniasinkam S, Díaz-Zuccarini V. Computational tools for clinical support: a multi-scale compliant model for haemodynamic simulations in an aortic dissection based on multi-modal imaging data. J R Soc Interface 2017;14.

[12] Dillon-Murphy D, Noorani A, Nordsletten D, Figueroa CA. Multi-modality image-based computational analysis of haemodynamics in aortic dissection. Biomech Model Mechanobiol 2016;15:857–76. doi:10.1007/s10237-015-0729-2.

[13] Cheng Z, Wood NB, Gibbs RGJ, Xu XY. Geometric and Flow Features of Type B Aortic Dissection: Initial Findings and Comparison of Medically Treated and Stented Cases. Ann Biomed Eng 2014;43:177–89. doi:10.1007/s10439-014-1075-8.

[14] Alimohammadi M, Agu O, Balabani S, Díaz-Zuccarini V. Development of a patient-specific simulation tool to analyse aortic dissections: Assessment of mixed patient-specific flow and pressure boundary conditions. Med Eng Phys 2014;36:275–84. doi:10.1016/j.medengphy.2013.11.003.

[15] Shang EK, Nathan DP, Fairman RM, Bavaria JE, Gorman RC, Gorman JH, et al. Use of computational fluid dynamics studies in predicting aneurysmal degeneration of acute type B aortic dissections. J Vasc Surg 2015;62:279–84. doi:10.1016/J.JVS.2015.02.048.

[16] Xu H, Li Z, Dong H, Zhang Y, Wei J, Watton PN, et al. Hemodynamic parameters that may predict false-lumen growth in type-B aortic dissection after endovascular repair: A preliminary study on long-term multiple follow-ups. Med Eng Phys 2017;7:35–0. doi:10.1016/j.medengphy.2017.08.011.

[17] Alimohammadi M, Sherwood JM, Karimpour M, Agu O, Balabani S, Díaz-Zuccarini V. Aortic dissection simulation models for clinical support: fluid-structure interaction vs. rigid wall models. Biomed Eng Online 2015;14:34. doi:10.1186/s12938-015-0032-6.

[18] Bonfanti M, Balabani S, Alimohammadi M, Agu O, Homer-Vanniasinkam S, Díaz-Zuccarini V. A simplified method to account for wall motion in patient-specific blood flow simulations of aortic dissection: Comparison with fluid-structure interaction. Med Eng Phys 2018;58:72–9. doi:doi:10.1016/j.medengphy.2018.04.014.

[19] Chen HY, Peelukhana S V., Berwick ZC, Kratzberg J, Krieger JF, Roeder B, et al. Editor’s Choice-Fluid-Structure Interaction Simulations of Aortic Dissection with Bench Validation. Eur J Vasc Endovasc Surg 2016;52:589–95. doi:10.1016/j.ejvs.2016.07.006.

[20] Crawford TC, Beaulieu RJ, Ehlert BA, Ratchford E V, Black JH, III. Malperfusion syndromes in aortic dissections. Vasc Med 2016;21:264–73. doi:10.1177/1358863X15625371.

[21] Gijsen FJH, van de Vosse FN, Janssen JD. The influence of the non-Newtonian properties of blood on the flow in large arteries: steady flow in a carotid bifurcation model. J Biomech 1999;32:601–8. doi:10.1016/S0021-9290(99)00015-9.

[22] Peacock J, Jones T, Tock C, Lutz R. The onset of turbulence in physiological pulsatile flow in a straight tube. Exp Fluids 1998;24:1–9. doi:10.1007/s003480050144.

[23] Alastruey J, Xiao N, Fok H, Schaeffter T, Figueroa CA, Parker KHKH, et al. On the impact of modelling assumptions in multi-scale, subject-specific models of aortic haemodynamics. J R Soc Interface 2016;13:20160073. doi:10.1098/rsif.2016.0073.

[24] Saouti N, Marcus JT, Noordegraaf AV, Westerhof N. Aortic function quantified: the heart’s essential cushion. J Appl Physiol 2012;113:1285–91. doi:10.1152/japplphysiol.00432.2012.

[25] Zamir M, Sinclair P, Wonnacott TH. Relation between diameter and flow in major branches of the arch of the aorta. J Biomech 1992;25:1303–10.

[26] Nakamura T, Moriyasu F, Ban N, Nishida O, Tamada T, Kawasaki T, et al. Quantitative measurement of abdominal arterial blood flow using image-directed Doppler ultrasonography: Superior mesenteric, splenic, and common hepatic arterial blood flow in normal adults. J Clin Ultrasound 1989;17:261–8. doi:10.1002/jcu.1870170406.

[27] Hall JE. Guyton and Hall Textbook of Medical Physiology. 13 edition. Saunders; 2015.

[28] Westerhof N, Stergiopulos N, Noble MIM. Snapshots of Hemodynamics. Boston, MA: Springer US; 2010. doi:10.1007/978-1-4419-6363-5.

[29] Les AS, Shadden SC, Figueroa CA, Park JM, Tedesco MM, Herfkens RJ, et al. Quantification of hemodynamics in abdominal aortic aneurysms during rest and exercise using magnetic resonance imaging and computational fluid dynamics. Ann Biomed Eng 2010;38:1288–313. doi:10.1007/s10439-010-9949-x.

[30] Chemla D, Hébert J-L, Aptecar E, Mazoit J-X, Zamani K, Frank R, et al. Empirical estimates of mean aortic pressure: advantages, drawbacks and implications for pressure redundancy. Clin Sci (Lond) 2002;103:7–13. doi:10.1042/.

[31] ANSYS Inc. CFX-Solver Theory Guide. Release 17. 2016.

[32] Gallo D, De Santis G, Negri F, Tresoldi D, Ponzini R, Massai D, et al. On the use of in vivo measured flow rates as boundary conditions for image-based hemodynamic models of the human aorta: Implications for indicators of abnormal flow. Ann Biomed Eng 2012;40:729–41. doi:10.1007/s10439-011-0431-1.

[33] Ioannou C V., Stergiopulos N, Katsamouris AN, Startchik I, Kalangos A, Licker MJ, et al. Hemodynamics induced after acute reduction of proximal thoracic aorta compliance. Eur J Vasc Endovasc Surg 2003;26:195–204. doi:10.1053/ejvs.2002.1917.

[34] Reymond P, Merenda F, Perren F, Ru D. Validation of a One-Dimensional Model of the Systemic Arterial Tree. Am J Physiol - Hear Circ Physiol 2009;297:208–22. doi:10.1152/ajpheart.00037.2009.

[35] Chemla D, Hébert J-L, Aptecar E, Mazoit J-X, Zamani K, Frank R, et al. Empirical estimates of mean aortic pressure: advantages, drawbacks and implications for pressure redundancy. Clin Sci (Lond) 2002;103:7–13. doi:10.1042/.

[36] Romagnoli S, Ricci Z, Quattrone D, Tofani L, Tujjar O, Villa G, et al. Accuracy of invasive arterial pressure monitoring in cardiovascular patients: an observational study. Crit Care 2014;18:644. doi:10.1186/s13054-014-0644-4.

[37] Pirola S, Cheng Z, Jarral OA, O’Regan DP, Pepper JR, Athanasiou T, et al. On the choice of outlet boundary conditions for patient-specific analysis of aortic flow using computational fluid dynamics. J Biomech 2017;60:15–21. doi:10.1016/j.jbiomech.2017.06.005.

[38] Itu L, Neumann D, Mihalef V, Meister F, Kramer M, Gulsun M, et al. Non-invasive assessment of patient-specific aortic haemodynamics from four-dimensional flow MRI data. Interface Focus 2018;8:20170006.

[39] Xiao N, Alastruey J, Figueroa CA. A systematic comparison between 1-D and 3-D hemodynamics in compliant arterial models. Int j Numer Method Biomed Eng 2014;30:204–31. doi:10.1002/cnm.2598.

[40] Rudenick PA, Bijnens BH, García-Dorado D, Evangelista A. An in vitro phantom study on the influence of tear size and configuration on the hemodynamics of the lumina in chronic type B aortic dissections. J Vasc Surg 2013;57:464–474.e5. doi:10.1016/j.jvs.2012.07.008.

[41] Soudah E, Rudenick P, Bordone M, Bijnens B, García-Dorado D, Evangelista A, et al. Validation of numerical flow simulations against in vitro phantom measurements in different type B aortic dissection scenarios. Comput Methods Biomech Biomed Engin 2015;18:805–15. doi:10.1080/10255842.2013.847095.

